# MicroRNA-21 restrains myD88-dependent protective inflammation worsening *Staphylococcus aureus skin* infection

**DOI:** 10.64898/2026.06.01.729375

**Authors:** Ana Carolina Guerta Salina, Amondrea Blackman, Alexandra Ivo de Medeiros, Carlos Henrique Serezani

**Affiliations:** Department of Medicine, Division of Infectious Diseases, Vanderbilt University Medical Center, Nashville, Tennessee, USA; Department of Pathology, Microbiology, and Immunology, Vanderbilt University Medical Center, Nashville, Tennessee, USA; Vanderbilt Institute of Infection, Immunology and Inflammation, Vanderbilt University Medical Center, Nashville, Tennessee, USA; Vanderbilt Center for Immunobiology, Vanderbilt University Medical Center, Nashville, Tennessee, USA; Department of Biological Sciences, School of Pharmaceutical Sciences, São Paulo State University, Araraquara, São Paulo, Brazil

**Author notes:** Address correspondence: Vanderbilt University Medical Center, Division of Infectious Diseases, Department of Medicine, Nashville, TN, USA. 1161 21^st^ Avenue South, AA6111, MCN, Nashville, TN, 37232.

**Keywords:** MiR-21, Methicillin-Resistant *Staphylococcus aureus*, Macrophages, Host defense, Skin, Abscess, Myeloid-differentiation protein 88

## Abstract

*Staphylococcus aureus* skin infections are driven by tissue-resident and recruited immune cells. A proper immune response requires a tightly regulated balance between inflammatory responses and the prevention of tissue damage. MicroRNAs are key post-transcriptional regulators of immune responses and influence both pro- and anti-inflammatory pathways. We show that *S. aureus* skin infection elevates miR-21 levels in the skin, and applying a topical miR-21 antagomir improves bacterial clearance and resolution of skin infection. In MRSA-infected mice lacking miR-21 in myeloid cells (miR21^Δmyel^), lesions are smaller, bacterial load decreases, macrophage infiltration is reduced, and collagen around the abscess increases. MiR-21 helps shape the inflammatory environment by modulating mediators such as IL-1β, TNF, IL-33, and IL-10. Additionally, miR-21 deficiency increases MyD88 expression in infected skin, and blocking MyD88 actions abolishes the protective effects observed in miR21^Δmyel^ mice. Overall, the miR-21/MyD88 pathway is a key regulator of the resolution of inflammation and antibacterial immunity during *S. aureus* skin infections, highlighting miR-21 inhibition as a promising therapy for antibiotic-resistant *S. aureus* infections.

## Introduction

*Staphylococcus aureus* is a Gram-positive bacterium commonly found in the human skin microbiota and is associated with opportunistic infections, ranging from asymptomatic colonization to invasive bacteremia, with a mortality rate of 20-30% (Kern 2010; Kluytmans et al. 1997; Tong et al. 2015). Antibiotic resistance is a public health concern affecting a broad spectrum of microorganisms and multiple antibiotic classes, including methicillin-resistant *Staphylococcus aureus* (MRSA), which is associated with skin infections, pneumonia, endocarditis, osteomyelitis, and sepsis (Kong et al. 2016; Ali Alghamdi et al. 2023; Kobayashi et al. 2015).

Host responses to *S. aureus* and MRSA infections are coordinated by resident tissue cells and recruited immune cells at the site of infection. During skin infection, resident macrophages and keratinocytes recognize pathogens through pathogen-associated molecular patterns (PAMPs) via pattern recognition receptors (PRRs), including toll-like receptors (TLRs) and scavenger receptors (Kobayashi et al. 2015; Bitschar et al. 2017; Brandt, Putnam, et al. 2018a; Miller 2008). Activation of the PAMP/PRR pathway induces production of cytokines, chemokines, and lipid mediators that orchestrate a robust influx of monocytes and neutrophils into infected tissue (Soares et al. 2023).

Neutrophils are essential for early infection control. These immune cells are rapidly recruited following infection and mediate pathogen clearance through phagocytosis, production of reactive oxygen and nitrogen species, and release of neutrophil extracellular traps (Zhang et al. 2024). Conversely, macrophages not only contribute to bacterial clearance but also are responsible for dead cell elimination (a process called efferocytosis) and tissue remodeling processes that support (Rodríguez-Morales and Franklin 2023).

Abscess formation is a fundamental host-defense mechanism against *S. aureus* and is a highly organized inflammatory structure composed of fibrin, both viable and dead cells, live bacteria, and tissue debris. Proper abscess maturation is characterized by fibroblast proliferation and the accumulation of pro-resolving macrophages that drive the formation of a collagen-rich fibrotic capsule surrounding the abscess. This architectural organization is essential for efficient bacteria containment, clearance, and subsequent tissue repair (Kobayashi et al. 2015; Bitschar et al. 2017; Brandt, Putnam, et al. 2018).

Inflammatory responses triggered by *S. aureus* infection must be tightly regulated to ensure pathogen clearance while limiting tissue damage. MicroRNAs (miRNAs) are small non-coding RNAs (∼18-25 nucleotides) that regulate gene expression post-transcriptionally by binding to the 3’-untranslated region (3’-UTR) of the target gene’s mRNA, resulting in translational repression or mRNA degradation. Several studies have demonstrated that *S. aureus* infection modulates the expression of multiple miRNAs, including microRNA-21 (miR-21), both *in vitro* and i*n vivo* (Jingjing et al. 2017; Luoreng et al. 2018; Tanaka et al. 2017; Ramirez et al. 2018; de Kerckhove et al. 2018; Kelly et al. 2025).

MiR-21 acts endogenously and can also be released in extracellular vesicles from multiple cells and organs (Y. Li et al. 2024), and it is the most abundant miRNA in macrophages (Canfrán-Duque et al. 2017). Although bioinformatic analyses predict hundreds of potential miR-21 targets, only a limited number of these interactions have been functionally validated (Kertesz et al. 2007). Previous studies from our group identified miR-21 as a key regulator of macrophage polarization, as miR-21 deficiency promotes enrichment of M2-like macrophages while reducing M1-associated macrophages (Wang et al. 2015). Also, miR-21 expression is associated with positive regulation of the transcription factor nuclear factor-κB (NF-κB) (Ma et al. 2011). Furthermore, miR-21 inhibits the expression and actions of the myeloid-differentiation protein 88 (MyD88) signaling pathway, suppressing inflammatory cytokine production (Scuruchi et al. 2024) while promoting IL-10 production and pro-resolving programs (De Melo et al. 2021). MyD88 signaling is a central component of protective immunity against infections caused by various pathogens, including *S. aureus* (Feng et al. 2014).

The role of miR-21 in host defense against *S. aureus* skin infections is not fully understood. Our study shows that miR-21 levels increase quickly during the early stages of *S. aureus* infections. MiR-21 inhibits MyD88 signaling, influences macrophage infiltration, and reduces collagen deposition, which affects abscess formation. Mice lacking miR-21 in myeloid cells develop smaller skin lesions, have better bacterial control, and show improved abscess development, along with less macrophage infiltration and more collagen. Additionally, blocking the MyD88 pathway identified the miR-21/MyD88 axis as a key regulator of immune response and inflammation resolution in *S. aureus* skin infections. Overall, these results suggest miR-21 could be a promising target for treating MRSA skin infections.

## Results

Inhibition of miR-21 promotes the resolution of skin infection caused by *S. aureus*.

Different bacterial infections are associated with upregulation of miR-21 in both *in vitro* and *in vivo* settings (Kelly et al. 2025; Johnston et al. 2017; Hackett et al. 2020; P. T. Liu et al. 2012). We assessed whether *S. aureus* skin infection induces miR-21 expression. Wild-type (WT) mice were infected subcutaneously, and skin biopsies were collected at multiple time points post-infection. MiR-21 levels increased early in infection (12 h), peaked at 24 h, and declined at 48 h post-infection **(Fig. 1A)**. To determine the biological relevance of miR-21 upregulation during infection, WT mice were infected and treated with an ointment containing antagomir-21 immediately after infection and daily for 9 days.

**Fig. 1.**
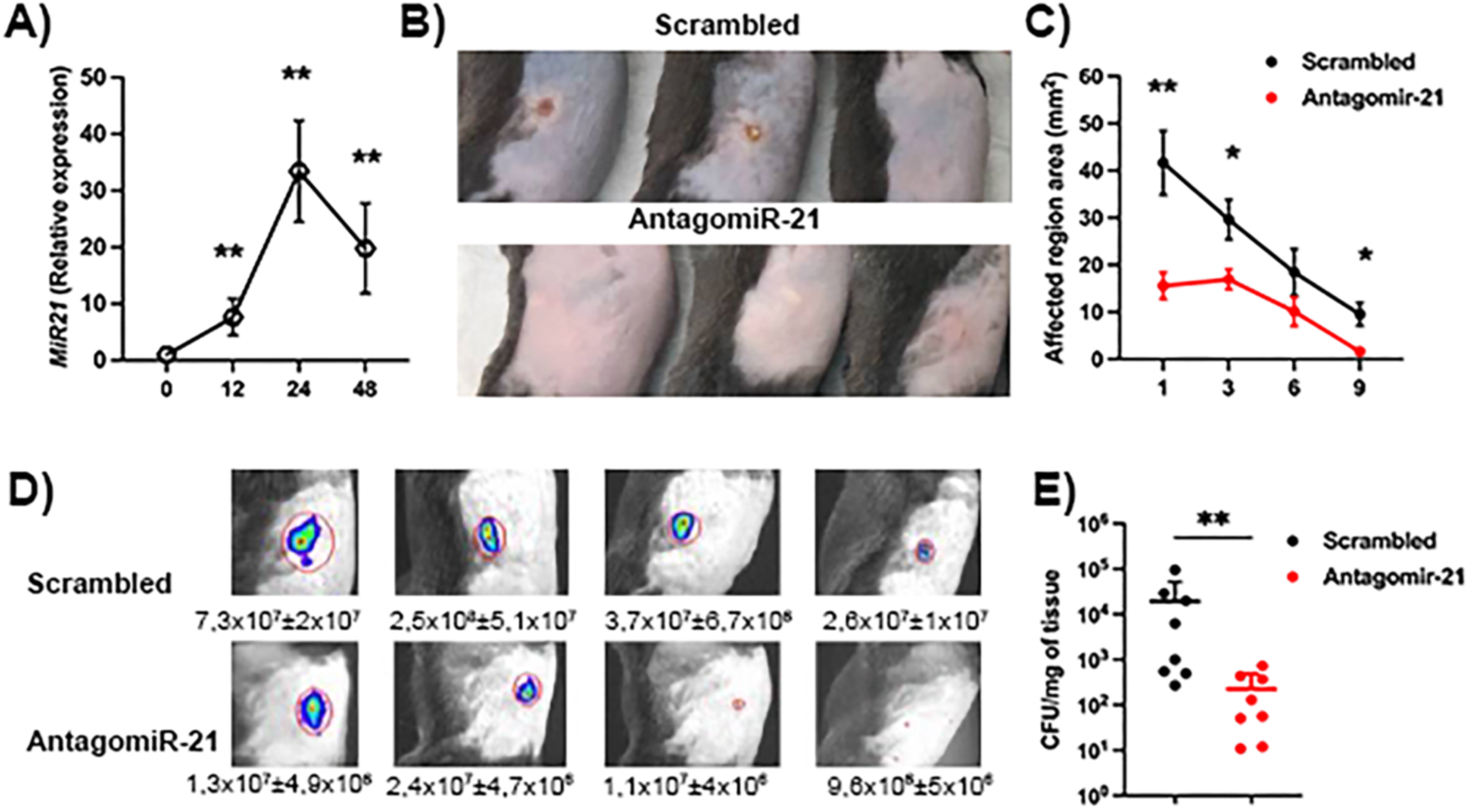
miR-21 inhibition enhances resolution of *S. aureus* skin infection. **A)** miR-21 expression was quantified in skin biopsies from WT mice infected with MRSA at time 0, 12 h, 24 h, and 48 h post-infection. Data are presented as mean ± SEM from three independent experiments. **p<0.005 *vs* t=0. **B-E)** WT mice were infected and treated daily with topical antagomir-21 or scrambled. **B)** Representative skin photographs collected at 24 h post-infection. **C)** Lesion size was monitored every other day for 9 days post-infection. **D)** Representative IVIS images and quantification of bioluminescence emitted by Biol-MRSA at days 1, 3, 6, and 9 post-infection. **E)** Bacterial burden in infected skin was quantified by CFU at day 9 post-infection. Data are presented as mean ± SD of six mice from a representative of two independent experiments. *p<0.05 and **p<0.005 *vs* antagomir control WT mice.

Notably, differences in lesion size were evident as early as 24 h post-infection. Mice treated with the miR-21 antagomir developed smaller skin lesions than those treated with a scrambled antagomir throughout the infection (**Fig. 1B-C**).

To monitor infection dynamics, we used a bioluminescent strain of *S. aureus* (biol-MRSA) to assess bacterial burden in individual animals throughout the infection. Mice treated with the miR-21 antagomir had lower bacterial loads than those treated with the scrambled antagomir (**Fig. 1D-E**). We confirmed this reduction by quantifying colony-forming units (CFUs) from skin biopsies collected on day 9 post-infection (**Fig. 1F**). Collectively, these data demonstrate that topical miR-21 inhibition enhances host immune responses during *S. aureus* skin infection.

### Macrophage-derived miR-21 expression impairs host responses against *S. aureus*

Next, we aimed to determine whether the effects of miR-21 are specific to myeloid cells. We infected miR-21 myeloid-deficient mice (miR-21^Δmyel^) and miR-21^flox/flox^ mice (control) and evaluated skin lesion size and bacterial burden.

MiR-21^Δmyel^ mice developed significantly smaller skin lesions throughout the infection compared with miR-21^flox/flox^ mice (**Fig. 2A-B**). This phenotype was correlated with reduced bacterial burden in miR-21^Δmyel^ mice at days 1, 3, and 6 post-infection in comparison to miR-21^flox/flox^ mice (**Fig. 2C**). Consistent with these findings, miR-21^Δmyel^ mice exhibited reduced CFUs in the skin at 24 h post-infection compared with control animals (**Fig. 2D**).

**Fig. 2.**
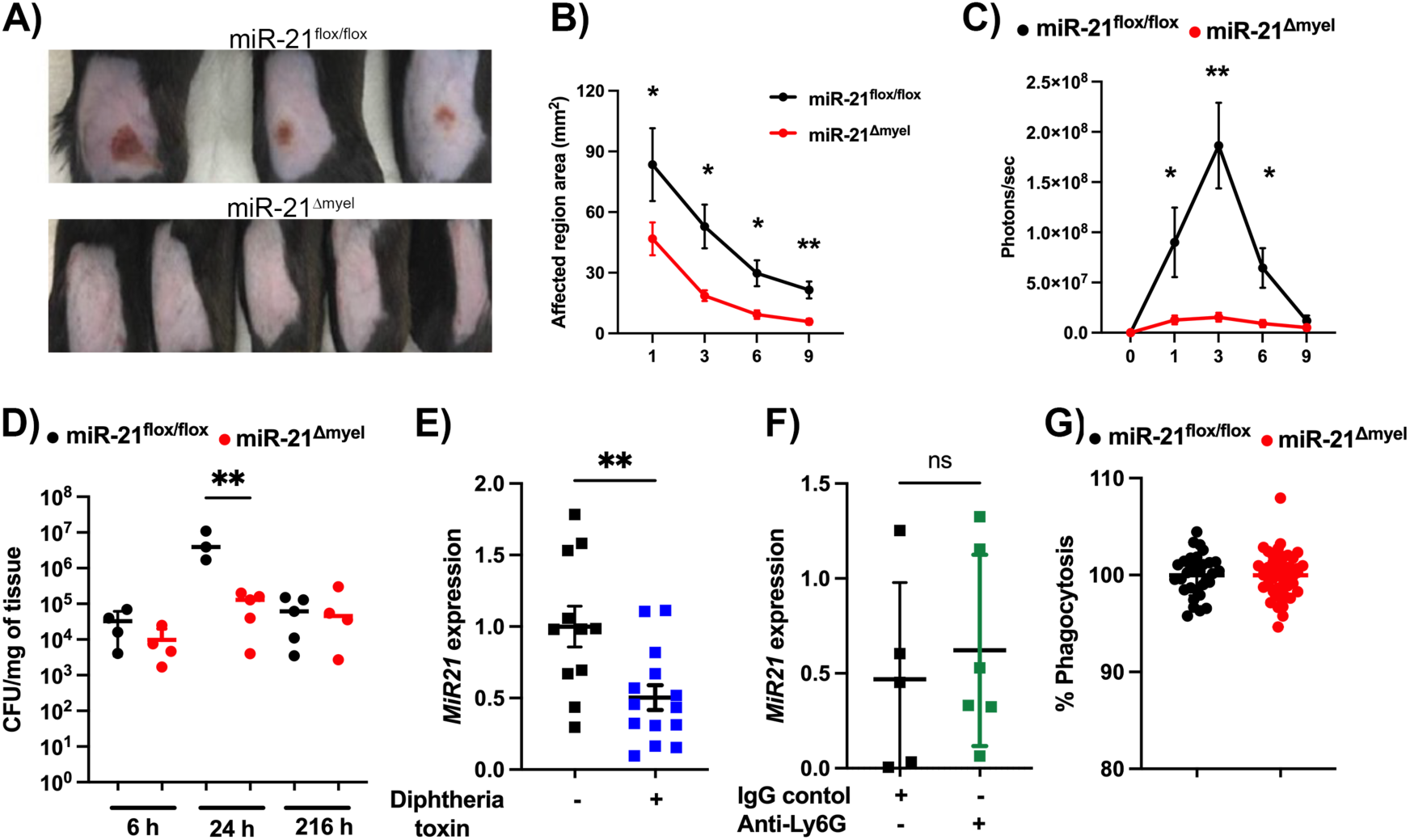
Myeloid-specific miR-21 deficiency promotes host defense. **A-D)** miR21^Δmyel^ or miR21^flox/flox^ mice were infected with Biol-MRSA and monitored throughout the infection. **A)** Representative skin photographs collected at 24 hours post-infection. **B)** Skin lesion size measured every other day for 9 days post-infection. **C)** Quantification of bioluminescence emitted by Biol-MRSA during infection using IVIS. Results are expressed as photons/second. **D)** Bacterial burden in infected skin quantified by CFU evaluation at 6h, day 1, and day 9 post-infection. Data are presented as mean ± SD of five mice from a representative of three independent experiments. *p<0.05 and **p<0.005 vs miR21^flox/flox^ mice. **E)** miR-21 expression was quantified in skin biopsies from MMDTR mice treated with diphtheria toxin (DT) or PBS at 24 h following MRSA infection, as described in Materials and Methods. Data are presented as mean ± SEM from two independent experiments. **p<0.005 vs MMDTR treated with PBS. **F)** miR-21 expression was quantified in skin biopsies from WT mice treated with anti-Ly6G antibody, as described in Materials and Methods, at 24 h following MRSA infection. Data are presented as mean ± SEM from two independent experiments. **G)** Peritoneal macrophages isolated from miR-21^Δmyel^ or miR-21^flox/flox^ mice were infected with GFP-expressing MRSA (MOI of 10), and microbial uptake was evaluated after 1 h. Data are expressed as the percentage of phagocytosis and presented as mean ± SEM from three independent experiments.

Macrophages and neutrophils are essential for clearing *S. aureus* during an infection (Kobayashi et al. 2015). To assess the individual contributions of these cells to early responses against *S. aureus* under conditions of reduced miR-21 expression, we selectively depleted miR-21 in macrophages and neutrophils. For macrophage depletion, we used MMDTR mice treated with diphtheria toxin prior to infection, thereby reducing miR-21 expression in the skin **(Fig. 2E)**. In parallel, we assessed the contribution of neutrophils to miR-21 expression using an anti-Ly6G antibody. Neutrophil depletion did not alter skin miR-21 expression **(Fig. 2F)**. We further confirmed that deleting miR-21 in neutrophils does not alter host responses to *S. aureus* infection in an *in vivo* model. Neutrophil miR-21-deficient (miR-21^ΔLy6G^) mice exhibited lesion sizes comparable to those of control animals throughout the course of infection, with no differences in bacterial burden in skin biopsies at day 9 post-infection **(Fig. S. 1A-B)**.

Subsequently, we examined whether miR-21 expression affects macrophages’ ability to ingest *S. aureus*. Macrophages from miR-21^Δmyel^ mice exhibited phagocytic capacity similar to macrophages from miR-21^flox/flox^ mice (**Fig. 2G**). Collectively, these findings indicate that actions of macrophage-derived miR-21 contribute to impaired host defense during *S. aureus* skin infection.

### miR-21 shapes the balance between pro- and anti-inflammatory cytokines during infection

Recognition of *S. aureus* by tissue-resident cells and recruited immune cells induces the production of cytokines that drive infection pathogenesis (Cho et al. 2012; Olaru and Jensen 2010; Puel et al. 2008; König et al. 1995; Cho et al. 2010). MiR-21^Δmyel^ mice exhibited an altered profile after infection, with increased levels of pro-inflammatory cytokines, including IL-1β, TNF, and IL-33, and reduced production of IL-6 and IL-1α in skin biopsies collected at day 1 post-infection **(Fig. 3A-E)**.

**Fig. 3.**
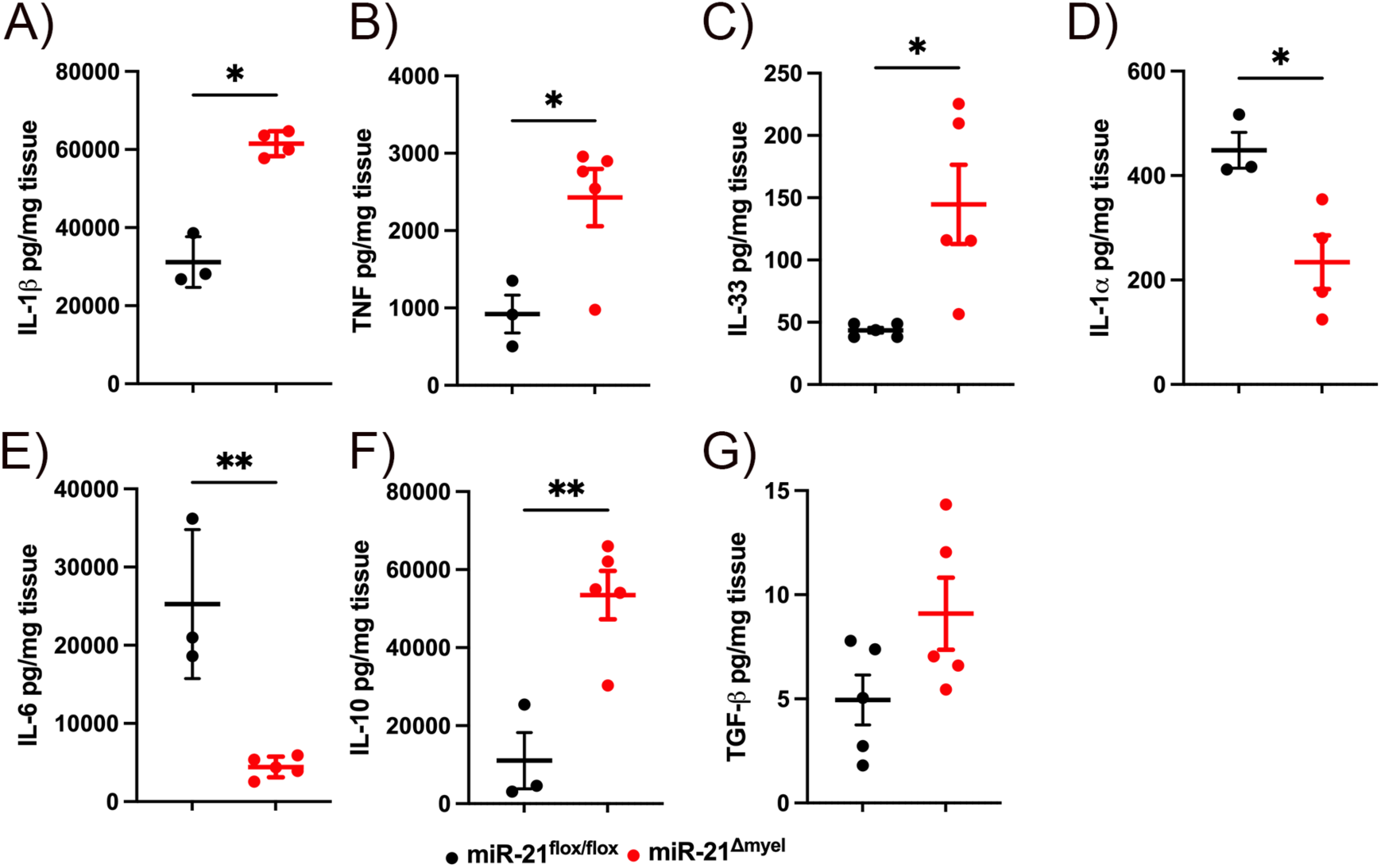
miR-21 controls the cytokine production during *S. aureus* infection. miR21^Δmyel^ or miR21^flox/flox^ mice were infected with MRSA, and skin biopsies were collected at 24 h post-infection for cytokine quantification by multiplex bead-based array assay or ELISA, as described in Materials and Methods. **A)** IL-1β, **B)** TNF, **C)** IL-33, **D)** IL-1α, **E)** IL-6, **F)** IL-10, and **G)** TGF-β. Data are presented as mean ± SD of at least 3 animals per group and are representative of three independent experiments. *p<0.05 and **p<0.005 *vs* miR21^flox/flox^ mice.

Interestingly, anti-inflammatory mediators were also affected, with miR-21^Δmyel^ mice exhibiting higher IL-10 levels and a trend toward increased TGF-β production compared with control animals (**Fig. 3F-G**). These results suggest that miR-21 plays a role in balancing pro- and anti-inflammatory cytokine responses during MRSA skin infection.

### Myeloid miR-21 deficiency improves abscess formation during *S. aureus* skin infection

We and others have previously shown that abscess formation is a critical host defense mechanism that promotes bacterial containment and clearance. Here, we observed that miR-21^Δmyel^ mice developed highly organized, spatially confined abscesses, characterized by a more clearly defined capsule and a more efficient containment of infiltrating cells at the infectious site compared with control animals (**Fig. 4A**).

**Fig 4.**
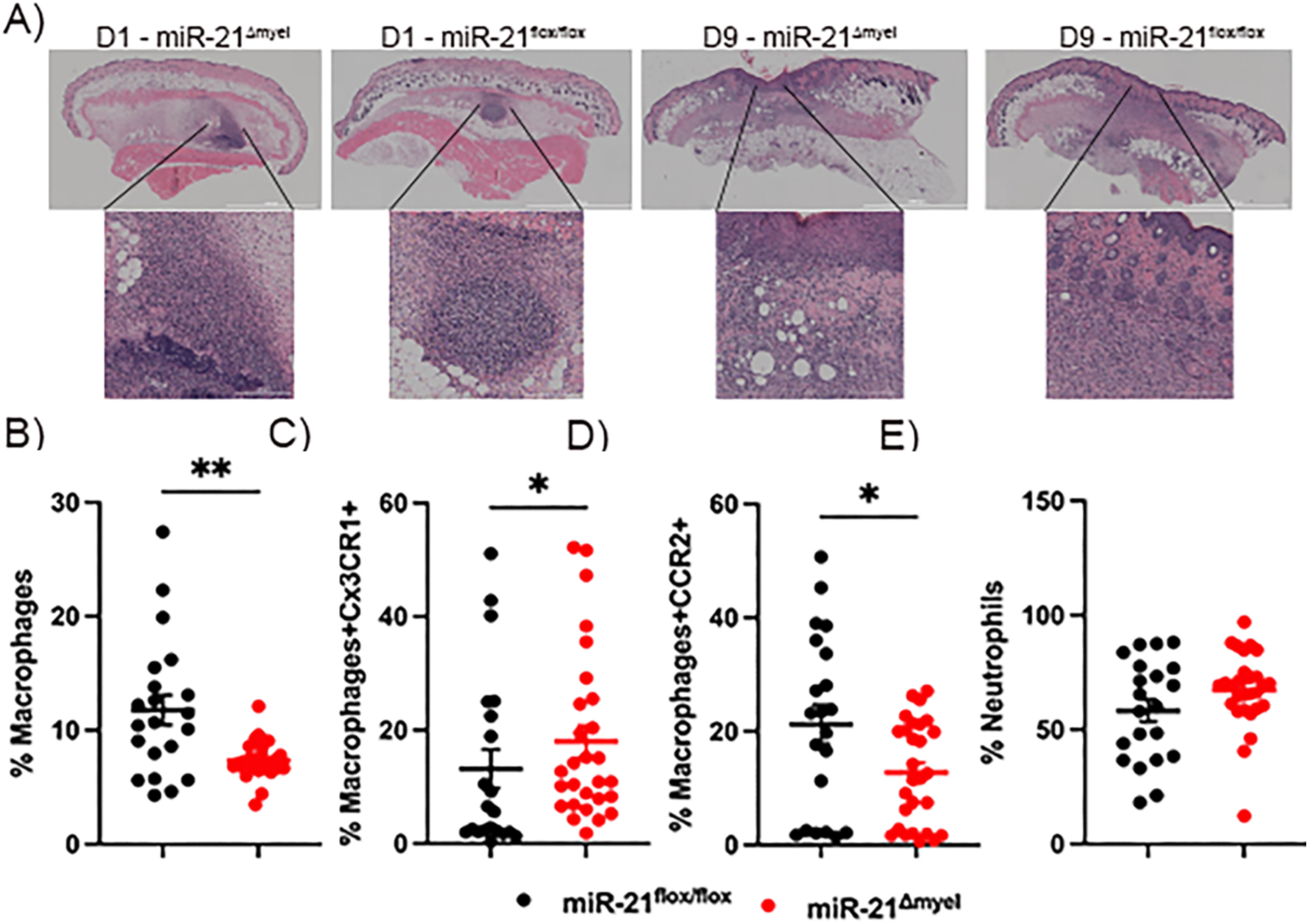
miR-21 deficiency mediates organized abscess formation and reduces the macrophage infiltration. miR21^Δmyel^ or miR21^flox/flox^ mice were infected with MRSA, and skin biopsies were collected at 24 h post-infection. **A)** Representative H&E staining of infected skin sections, magnification 20x. **B-E)** Skin tissue was digested into single-cell suspensions, as described in Materials and Methods, and immune cell populations were characterized by flow cytometry. **B)** Percentage of total macrophages (gated on total cells/ single cells/ live cells/ CD64+). **C)** Percentage of resident macrophages expressing CX3CR1+ (gated on total cells/ single cells/ live cells/ CD64+ CX3CR1+). **D)** Percentage of infiltrating macrophages expressing CCR2+ (gated on total cells/ single cells/ live cells/ CD64+ CCR2+). **E)** Percentage of neutrophils (gated on total cells/ single cells/ live cells/ Ly6G+). Data are presented as mean ± SEM from three independent experiments. *p<0.05 and **p<0.005 *vs* miR21^flox/flox^.

We observed that miR-21^Δmyel^ mice exhibited reduced macrophage accumulation in the tissue at 24 h post-infection compared with miR-21^flox/flox^ mice (**Fig. 4B**). Notably, this reduction was accompanied by a shift in macrophage composition, with miR-21^Δmyel^ mice displaying a higher proportion of resident macrophages (CD64+CX3CR1+ cells), whereas infiltrating macrophages (CD64+CCR2+ cells) predominated in control tissues (**Fig. 4C-D**). In contrast, the proportion of recruited neutrophils was comparable between groups, indicating that neutrophil recruitment was not affected (**Fig. 4E**).

The formation of a collagen-rich fibrotic capsule is essential for proper abscess development and maturation (Roy et al. 2020). We measured collagen deposition in skin biopsies from miR-21^Δmyel^ and control mice using trichrome blue staining. MiR-21^Δmyel^ mice showed significantly increased collagen levels compared to control (**Fig. 5A-B**). Supporting this, *Col1a3* expression was also higher in the skin of miR-21^Δmyel^ mice 24 h after infection (**Fig. 5C**).

**Fig. 5.**
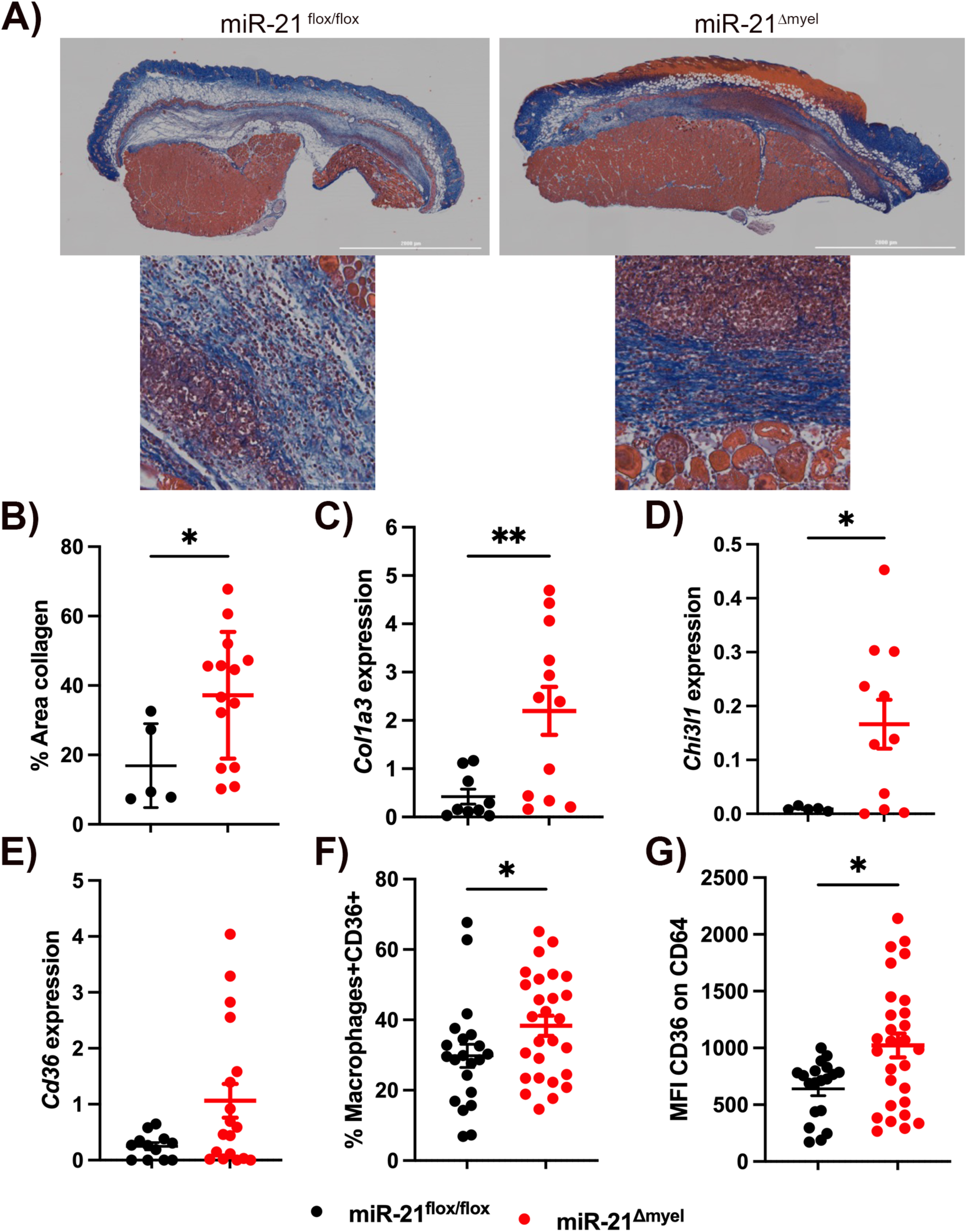
miR-21 deficiency enhances collagen deposition and pro-resolving macrophages during *S. aureus* skin infection. miR21^Δmyel^ or miR21^flox/flox^ mice were infected with MRSA, and skin biopsies were collected at 24 h post-infection. **A)** Representative Masson’s trichrome blue staining of infected skin sections at 20x and 40x magnification. **B)** Quantification of collagen deposition from Masson’s trichrome blue sections using ImageJ. Results are expressed as the percentage of collagen-positive area relative to total tissue area. **C-E)** Skin biopsies were processed for RT-qPCR analysis to determine expression of **C)** *Col1a3,* **D)** *Chi3l1,* and **E)** *Cd36*. **F-G)** Skin tissue was digested into a single-cell suspension, as described in Materials and Methods, and immune cell populations were characterized by flow cytometry. **F)** Percentage of CD36+ macrophages (gated on total cells/single cells/live cells/CD64+ CD36+). **G)** Mean Fluorescence Intensity (MFI) of CD36 expression in macrophages. Data are presented as mean ± SEM from three independent experiments. *p<0.05 and **p<0.005 *vs* miR21^flox/flox^.

Interestingly, elevated *Col1a3* expression was also detected in the skin of mice treated with antagomir-21 at 9 days post-infection (**Fig. S. 3A**). Furthermore, we observed increased expression of pro-resolving genes, including *Chil1l3* and *Cd36,* in tissues from mice topically treated with antagomir-21 (**Fig. S. 3B-C**). Consistent with these findings, miR-21^Δmyel^ mice exhibited increased *Chi3l1* expression (**Fig. 5D**) and a higher proportion of macrophages expressing CD36 (CD64+CD36+ cells) at 24h post-infection (**Fig. 5F - G**).

Collectively, these findings indicate that miR-21 hinders pro-resolving macrophage responses and promotes excessive recruitment of circulating macrophages, while disrupts abscess organization and maturation during *S. aureus* skin infection.

### miR-21 suppresses MyD88-dependent host defense signaling

Studies have demonstrated that MyD88 is essential for effective host defense against *S. aureus* infection by promoting immune cell recruitment and the production of inflammatory cytokines (Takeuchi et al. 2000; Miller et al. 2006). We detected increased *Myd88* expression in the skin of MRSA-infected miR-21^Δmyel^ mice compared with infected miR-21^flox/flox^ mice (**Fig. 6A**). Moreover, macrophages isolated from the infected tissue of miR-21^Δmyel^ mice showed higher MyD88 expression (CD64+MyD88+ cells) (**Fig. 6B-C**).

**Fig. 6.**
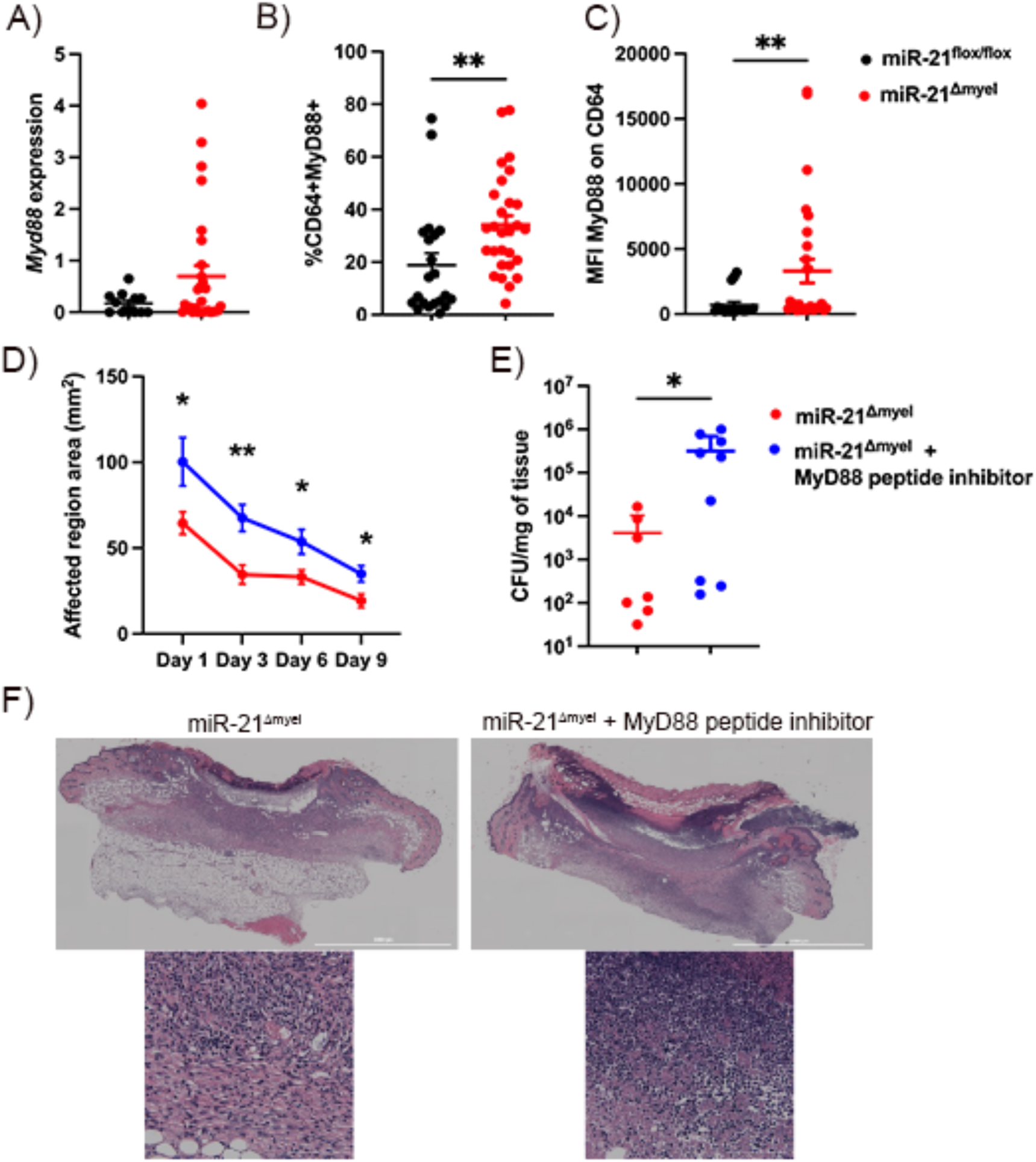
miR-21 suppresses MyD88-dependent host defense during S. aureus skin infection. **A-C)** MiR21Δmyel or miR21flox/flox mice were infected with MRSA, and biopsies were collected at 24 h post-infection. **A)** MyD88 expression in skin biopsies was determined by RT-qPCR. **B-C)** Skin tissue was digested into a single-cell suspension, as described in Materials and Methods, and the immune cell population was characterized by flow cytometry. **B)** Percentage of MyD88+ macrophages (gated on total cells/single cells/live cells/CD64+ MyD88+). **C)** Mean Fluorescence Intensity (MFI) of MyD88 expression in macrophages. Data are presented as mean ± SEM from three independent experiments. **p<0.005 vs miR21^flox/flox^. **D-F)** MiR21Δmyel mice were infected with MRSA and treated daily with a MyD88 peptide inhibitor or peptide control by intraperitoneal injection. **D)** Skin lesion size was measured every other day for 9 days post-infection. **E)** Bacterial burden in infected skin was quantified by CFU at day 9 post-skin infection. **F)** Representative H&E staining of infected skin sections collected at day 9 post-infection, magnification 20x and 40x, respectively. Data are presented as mean ± SD of at least six lesions from a representative of three independent experiments. *p<0.05 and **p<0.005 vs miR21^Δmyel^ mice.

To assess whether miR-21’s suppression of MyD88-dependent signaling plays a functional role in host defense against *S. aureus* infection, miR-21^Δmyel^ mice were administered with a MyD88 inhibitory peptide or control peptide throughout the infection. Disease progression was monitored by measuring skin lesion size over time, and bacterial loads were quantified at day 9 post-infection. Pharmacological blocking of MyD88 abolished the protective effect observed in miR-21^Δmyel^ mice, leading to significantly larger skin lesions compared to peptide-treated mice (**Fig. 6D**). This was also associated with an increased bacterial burden at day 9 post-infection (**Fig. 6E**). Histopathological analysis revealed that, although organized abscesses were absent at this stage, skin biopsies from mice treated with the MyD88 inhibitor peptide displayed inflammatory infiltrates and delayed tissue healing versus control (**Fig. 6F**). Overall, these results confirm the miR-21/MyD88 pathway as a key regulator of antibacterial responses and of inflammatory resolution during *S. aureus* skin infection.

## Discussion

MicroRNAs tightly regulate all stages of infection, from the initial host-pathogen interaction to tissue recovery, by controlling the post-transcriptional expression of up to 60% of protein-coding genes (Friedman et al. 2009). Many of these gene targets are involved in the innate and adaptive immune pathways that coordinate bacterial sensing and clearance, apoptotic cell removal, and tissue repair programs required for wound healing (Tanaka et al. 2017; de Kerckhove et al. 2018; Drury et al. 2017). Previous studies have demonstrated that distinct microRNAs regulate host responses during *S. aureus* infection across multiple tissues (Jingjing et al. 2017; Luoreng et al. 2018; Tanaka et al. 2017; Ramirez et al. 2018; de Kerckhove et al. 2018). Here, we demonstrate that *S. aureus* infection induces miR-21 expression in the skin and that loss of miR-21 enhances bacterial clearance and reduces lesion severity. Mechanistically, miR-21 deficiency inhibits the accumulation of infiltrating macrophages and promotes the formation of highly organized abscesses with increased collagen deposition, and enhancing bacterial containment. Furthermore, pharmacological inhibition of MyD88 signaling abolishes the protective phenotype observed in miR-21-deficient mice, identifying the miR-21/MyD88 axis as a critical regulator of antibacterial immunity and inflammatory resolution during *S. aureus* skin infection.

The first hours of infection are crucial for effective bacteria containment, a process coordinated by rapid activation of multiple immune pathways and molecules. We observed that MRSA skin infection rapidly induces miR-21 expression. Corroborating our findings, miR-21 upregulation has been reported across a broad spectrum of infectious settings, incluinding infections caused by *Mycobacterium leprae* (P. T. Liu et al. 2012), *Leishmania donovani* (Varikuti et al. 2021), Brucella spp (Rezaeepoor et al. 2024), *S. aureus* (Kelly et al. 2025), and hepatitis C virus (H.-C. Li et al. 2022).

In the skin, the early inflammatory response to *S. aureus* culminates in abscess formation, with lesion and abscess size mirroring the extent of local tissue damage and bacterial burden within the infected tissue (Kobayashi et al. 2015). We observed that both pharmacological blockade of miR-21 and selective depletion of this microRNA in myeloid cells enhance host control of infection, limiting bacterial burden and lesion progression.

Recognition of *S. aureus* by keratinocytes and macrophages triggers the rapid production of chemotactic mediators and inflammatory cytokines, promoting robust neutrophil recruitment and establishment of an inflammatory microenvironment (Abtin et al. 2014; Minegishi et al. 2009). This response must be tightly coordinated, balancing pro-inflammatory mediators that drive immune cell activation, and anti-inflammatory pathways that limit excessive tissue damage and promote wound healing (Riaz et al. 2025; Dai and Chen, n.d.). In this context, skin biopsies from *S. aureus*-infected miR-21^Δmyel^ mice displayed elevated levels of IL-1β, IL-33, and TNF, as well as increased IL-10 and TGF-β production, compared to miR-21^flox/flox^ mice. TNF produced by keratinocytes, mast cells, and dermal macrophages has been shown to promote *S. aureus* clearance by enhancing the recruitment of immune cells to sites of infection (Aufiero et al. 2007; C. Liu et al. 2018). Likewise, IL-1β generated during *S. aureus* infection is critical for recruitment of neutrophils and effective host defense (Cho et al. 2012; Miller et al. 2006),(Putnam et al. 2019). Therefore, the higher levels of IL-1β and TNF seen in miR-21^Δmyel^ mice likely contributed to the improved bacterial clearance noted in our experimental setup.

We also observed increased IL-10 and TGF-β levels in infected skin miR21^Δmyel^ mice, suggesting simultaneous activation of regulatory pathways that restrain excessive inflammation. Stenzel and colleagues demonstrated that IL-10-deficient mice infected with *S. aureus* exhibit increased susceptibility in a brain abscess model characterized by necrotizing vasculitis, hemorrhage, and severe edema formation (Stenzel et al. 2005). Although Sheedy and collaborators previously reported that miR-21 enhances IL-10 production by repressing the IL-10 inhibitor PDCD4 (Sheedy et al. 2010), it is important to note that miR-21 induction in the study was induced by LPS stimulation. In contrast, our experimental model employed a live pathogen, introducing a substantially more complex response. Moreover, LPS is a component of Gram-negative bacteria, whereas *S. aureus* is a Gram-positive pathogen.

Abscess formation is a critical host-defense mechanism during *S. aureus* infection and can occur in multiple tissues, including the skin (Abtin et al. 2014; Stenzel et al. 2005), kidney (Cheng et al. 2009), and brain (Kielian et al. 2004). Although abscesses exhibit tissue-specific characteristics depending on the affected organ, their structural organization and maturation processes are similar (Kobayashi et al. 2015). The formation of a fibrotic capsule by macrophages and fibroblasts prevents bacterial dissemination (Brandt, Putnam, et al. 2018) while promoting neutrophil accumulation at the site of infection, contributing to microbial containment (Pasparakis et al. 2014). Our results demonstrated that miR-21^Δmyel^ mice exhibited improved abscess organization during *S. aureus* skin infection, characterized by enhanced collagen-rich capsule formation and limited macrophage infiltration into the infectious core. These findings suggest that miR-21 impairs the establishment of a protective tissue-containment program during *S. aureus* skin infection by limiting abscess capsule formation, compromising bacterial confinement and abscess maturation.

Macrophages exhibit notable functional plasticity, enabling them to dynamically adapt to the microenvironment. While classically activated macrophages are associated with pro-inflammatory and antimicrobial activities, M2-like macrophages are known for producing anti-inflammatory mediators and for their role in tissue remodeling, including collagen deposition and angiogenesis (Y.-C. Liu et al. 2014). In this study, we observed an enrichment of transcriptional programs tied to pro-resolving and tissue-remodeling macrophages. Consistent with this, Aldrich and Kielian demonstrated that *S. aureus* infection promotes the accumulation of reparative macrophages in the brain abscess wall, where these cells help organize and mature the abscess (Aldrich and Kielian 2011). Moreover, miR-21 has been identified as a crucial regulator of macrophage reprogramming, with miR-21-deficient mice displaying an increase in tissue-remodeling macrophages and a reduction in inflammatory macrophage signatures (Wang et al. 2015)

MyD88 is a master adaptor protein required for signaling induced by most toll-like receptors, except TLR3, as well as IL-1 family cytokine receptors (Deguine and Barton 2014; Kawai and Akira 2010). Loss of MyD88 signaling severely compromises neutrophil recruitment and inflammatory cytokine production, increasing susceptibility to *S. aureus* infection (Takeuchi et al. 2000; Miller et al. 2006, 2007). Furthermore, previous studies have identified miR-21 as a negative regulator of the MyD88/ NF-κB signaling axis, restraining inflammatory cytokine production while promoting anti-inflammatory and pro-resolving transcriptional programs, including IL-10 expression (Sheedy et al. 2010; Feng et al. 2014; Wang et al. 2015). Consistent with this regulatory network, we observed increased MyD88 expression in the skin of miR21^Δmyel^ mice at 24 h post-infection. Skin biopsies collected from miR21^Δmyel^ mice displayed an increased frequency of macrophages expressing MyD88 during *S. aureus* skin infection. Notably, pharmacological blockade of this signaling axis via administration of a MyD88-inhibitory peptide abolished the protective phenotype observed in miR21^Δmyel^ mice, resulting in more severe skin lesions and increased bacterial burden. Our results also showed that even 9 days after infection, miR21^Δmyel^ mice treated with the MyD88-inhibitory peptide continued to exhibit inflammatory cell infiltration and impaired restoration of tissue homeostasis. Overall, these findings highlight miR-21 as an essential immunoregulatory checkpoint in *S. aureus* skin infection, where its expression inhibits MyD88-dependent innate immune responses. This suppression hampers bacterial containment, disrupts abscess formation, and ultimately worsens tissue damage.

Despite major advances in antimicrobial therapy, *S. aureus* skin infections remain a significant clinical challenge, particularly with multidrug-resistant strains, which are frequently associated with extensive tissue destruction, impaired wound resolution, and systemic dissemination (Kong et al. 2016; Ali Alghamdi et al. 2023). These limitations highlight the urgent need for host-directed therapeutic strategies that enhance antimicrobial immunity while preserving tissue integrity. In this context, we identify miR-21 as a negative regulator of protective host defense during *S. aureus* skin infection. Our findings suggest that miR-21 restrains MyD88-dependent inflammatory signaling, compromises abscess organization, and limits effective bacterial containment. Thus, inhibition of miR-21 with antagomiR-21 may offer a promising therapeutic strategy to restore antimicrobial inflammation, improve host control of *S. aureus*, and promote tissue repair.

## Materials and Methods

### Animals

Mice were maintained in accordance with NIH guidelines for the use of experimental mice, with approval from the Institutional Animal Care and Use Committee at VUMC (Protocol Number: #M1600215). miR-21^flox/flox^ mouse strain was generated by Dr. Ivan Mircea through Ozgene Ptv Ltd (Perth, Australia). miR-21 myeloid-deficient mice – miR-21^Δmyel^ (C57BL/6 background) - were generated by breeding the miR-21^flox/flox^ with LysM^cre^. C57BL/6 (WT) and MMDTR mice were purchased from Jackson Labs (Bar Harbor, ME, USA). MMDTR mice were generated by breeding the Csf1r^HBEGF/mCherry1Mnz/J^ with LysM^cre^ mice (C57BL/6 background), as previously reported(Schreiber et al. 2013). Neutrophil miR-21-deficient (miR-21^ΔLy6G^) mice were generated by breeding miR-21^flox/flox^ with Ly6g^tm2621(Cre,-tdTomato)Arte^ (C57BL/6 background). Male and female mice between 8 and 12 weeks of age were used in all experiments.

### Methicillin-Resistant *Staphylococcus aureus* Strain

The bioluminescent methicillin-resistant *Staphylococcus aureus* (NRS384 lux) strain was provided by Dr. Roger Plaut (Food and Drug Administration, Silver Spring, Maryland, USA) (Plaut et al. 2013). The GFP-expressing USA300 LAC strain was provided by Dr. William Nauseef (University of Iowa, Iowa City, Iowa) (Greenlee-Wacker et al. 2014). MRSA stocks were stored at -80 °C and cultivated in tryptic soy broth medium (TSB; BD Biosciences) as previously described(Dejani et al. 2016).

### MRSA skin infection and mice treatment

Mice were infected subcutaneously with approximately 5×10^6^ colony-forming units (CFU) of MRSA in 50 μL of phosphate-buffered saline (PBS) as previously described (Dejani et al. 2016). Skin lesion size was monitored daily, and the affected area was calculated by the standard equation for the area [area = (π/2) × length × width](Becker et al. 2014).

Topical ointments were prepared using 100% petroleum jelly (Vaseline) emulsified immediately before application with antagomir-21 (100 ng/g) (Integrated DNA Technologies - IDT) or scrambled-antagomir control (100 ng/g; IDT). Ointments were applied daily using a sterile cotton swab to fully cover the infected area.

MyD88 inhibitor peptide (TJ-M2010-5; MedChemExpress - MCE; Cat n° HY-139397) at 50mg/kg or Antennapedia control peptide (Novus; Cat n° NBP2-29334) at 0.1mg/kg were reconstituted in 100% corn oil (MCE; Cat n° HY-Y1888). Daily intraperitoneal treatments were administered at 200 μl per mouse throughout the study period.

### Skin biopsies, harvest, and bacterial load

Punch biopsies of skin (8 mm) were collected at different post-infection time points (ranging from 6 hours to 9 days) and used to determine bacterial counts, assess cytokine production, extract RNA, perform single-cell characterization, and conduct histological analyses (Novelli et al. 2000). For bacterial load, skin biopsies were collected and homogenized in TSB medium. Serial dilutions were plated on tryptic soy agar (TSA) medium (BD), and the plates were incubated at 37°C, 5% CO2 for 18h, and then CFU counts were performed. Bacterial burdens were normalized to tissue weight and calculated using the following equation: ((CFU/mL plated) * (dilution factor)) / (tissue weight in mg). Bacterial burdens in the skin are represented by CFU/mg of tissue (Dejani et al. 2016).

### RNA isolation and quantitative real-time RT-PCR

Skin biopsies were collected in RNAlater stabilization buffer. For quantification of *Myd88*, *Col1a3*, *Cd36,* and *Chil3l1* expression, tissues were homogenized in RLT lysis buffer, and total RNA was isolated using the QIAGEN RNeasy Mini Kit according to the manufacturer’s instructions (Qiagen Hilden; Cat n° 74104). cDNA was synthesized using the iScript cDNA Synthesis Kit (Bio-Rad; Cat n° 1708891).

To quantify miR-21, total microRNA was extracted with the Direct-zol RNA Miniprep Kit (Zymo Research; Cat n° R2050), and cDNA was synthesized using the Mir-XTM miRNA First-Strand Synthesis Kit (Takara, Cat n° 638315) per manufacturers’ instructions. In both instances, quantitative PCR (qPCR) was performed on a CFX96 Real-Time PCR Detection System (Bio-Rad Laboratories). All primers were sourced from IDT.

### *In vivo* imaging (IVIS)

Mice were anesthetized with isoflurane and imaged *in vivo* using an IVIS Spectrum/CT optical imaging system (PerkinElmer), as previously described(Brandt, Klopfenstein, et al. 2018). Serial longitudinal scans wereaquired throughout the MRSA skin infection. To quantify infection severity, a region of interest around the site of infection was defined, and the total flux (photons/second) recorded.

### MMDTR mice and macrophage ablation

To deplete resident skin macrophages, MMDTR mice were treated with 100 ng Diphtheria toxin (DT; Sigma-Aldrich; Cat n° D0564) or PBS vehicle control intraperitoneally daily for 3 consecutive days prior to MRSA skin infection. Skin biopsies were collected 24 h post-infection.

### Neutrophil depletion

Mice were treated intraperitoneally with 200 μg of anti-Ly6G-depleting antibody (clone 1A8; BioXcell; Cat n° BE0075-1) or the IgG isotype control (Cat n° BE0089). Antibodies were administered 18 h prior to MRSA skin infection as described (Brandt, Klopfenstein, et al. 2018). Skin biopsy samples were collected 24 h post-infection.

### Phagocytosis assays

Peritoneal macrophages (PM) from miR21^Δmyel^ or miR-21^flox/flox^ mice were harvested from the peritoneal cavities by lavage with ice-cold PBS and plated at 2 × 10^5^ cells per well in a black 96-well plate with clear bottoms (Thermo Fisher Scientific, Cat n° 1665305). Cells were allowed to adhere for 18 h in complete DMEM supplemented with 10% heat-inactivated FBS, 1% penicillin-streptomycin, and 1% L-glutamine (200 mM). PMs were infected with GFP-MRSA at a multiplicity of infection (MOI) of 10 (Brandt, Klopfenstein, et al. 2018). Following 1h of infection, wells were washed with warm PBS, and GFP fluorescence was measured. To determine the intensity of intracellular fluorescence, extracellular GFP was quenched with 500 μg/mL trypan blue, and the GFP fluorescence was quantified using a fluorimeter plate reader.

### Cytokine detection

Skin biopsies were collected and homogenized in TNE cell lysis buffer containing 1X protease & phosphatase inhibitor cocktail (Thermo Fisher Scientific; Cat n° 78442). Levels of IL-1α, IL-1β, IL-10, IL-6, and TNF-α were quantified using a multiplex bead-based array assay performed by Eve Technologies (Eve Technologies). IL-33 and TGF-β concentrations were determined by ELISA according to the manufactureŕs instructions (R&D Systems; Cat n° DY3626 and DY1679, respectively). Cytokine levels were normalized to tissue weight and data expressed as pg/mg of tissue.

### Histopathological analysis

Skin biopsies were fixed in 4% paraformaldehyde and embedded in paraffin. Tissue sections were stained with hematoxylin and eosin (H&E) to visualize abscesses and with Masson’s trichrome blue to detect collagen. Images were captured at 20x or 40x magnifications using an Agilent BioTek Cytation 7 multimode cell imaging reader (BioTek). Collagen deposition was quantified using ImageJ software v1.54g.

### Skin single-cell isolation and flow cytometry

Skin biopsy specimens were collected, minced, and digested in 3 mL of DMEM containing 2 mg/mL collagenase Type XI (Roche Diagnostics, Cat n° C7657) and 0.5mg/mL hyaluronidase (Sigma-Aldrich, Cat n° H3506) for 45 minutes at 37°C. Single-cell suspensions were incubated with CD16/32 Fc-blocking antibodies (Biolegend; Cat n°101310; clone 93) to prevent non-specific antibody binding, followed by staining with the viability dye (Ghost Dye^TM^ Violet 510, Cytak, Cat n° 13-0870-T100) for 30 minutes. Next, cells were fixed and permeabilized using the BD Cytofix/Cytoperm Kit (Cat n° 554714) and stained with fluorescently labeled antibodies for 30 minutes. The following antibodies were used: CD64-PE/Cy7 (Biolegend; Cat n° 139314; clone X54-5/7.1), Ly6G-APC (Biolegend; Cat n° 127614; clone 1A8), CX3CR1-PerCP/Cy5.5 (Biolegend; Cat n° 149010; clone SA011F11), CCR2-FITC (Biolegend; Cat n° 141703; clone C068C2), CD36-PE (Biolegend; Cat n° 102605; clone HM36), MyD88 (Abcam; Cat n° ab2064), and goat anti-rabbit IgG-Alexa Fluor^TM^ 647 (Invitrogen; Cat n° A21245). Analyses were performed using FlowJo v.10 software.

### Statistical Analysis

Data are presented as mean ± SEM for pooled biological experiments or mean ± SD for representative experiments. Statistical analyses were performed using GraphPad Prism v10.5.0 software. Comparisons between two experimental groups were performed using a two-tailed Student’s *t-* test, whereas comparisons among three or more experimental groups were analyzed using one-way ANOVA or two-way ANOVA followed by Tukey’s multiple-comparison test, as appropriate. P values < 0.05 were considered statistically significant.

## Acknowledgments

We would like to thank the Serezani laboratory for their input. We also extend our thanks to Bethany Moore (University of Michigan, Ann Arbor, MI, USA) for providing the MRSA USA300 LAC strain and to Roger Plaut (Food and Drug Administration, Silver Spring, MD, USA) for supplying the bioluminescent USA300 (NRS384 lux) MRSA strain. This work was supported by NIH grants: 1R01AI180777-01, R01DK122147 (to CHS). ACGS was funded by 5TL1TR002244-09 and Fundação de Amparo à Pesquisa do Estado de São Paulo (FAPESP; grants: # 2017/04786-0 and # 2018/01622-9). AIM was funded by Conselho Nacional de Desenvolvimento Científico e Tecnológico 308705/2022-0-CNPQ-PQ

We also acknowledge: 1) Vanderbilt Flow Cytometry Shared Resource, supported by the Vanderbilt Ingram Cancer Center (P30 CA068485) and the Vanderbilt Digestive Disease Research Center (DK058404); 2) Translational Pathology Shared Resource supported by NCI/NIH Cancer Center Support Grant P30CA068485.

## Author Contribution

ACGS: Conceptualization, Data curation, Formal analysis, Investigation, Methodology, Writing – original draft.

AB: Data curation, Investigation, Methodology.

AIM: Conceptualization.

CHS: Conceptualization, Formal analysis, Funding acquisition, Project administration, Resources, Supervision, Writing – original draft.

All authors read and approved the final manuscript.

## Declaration of AI/LLM

The authors state no AI/LLM use.

## FIGURES

**Fig. Supp 1.**
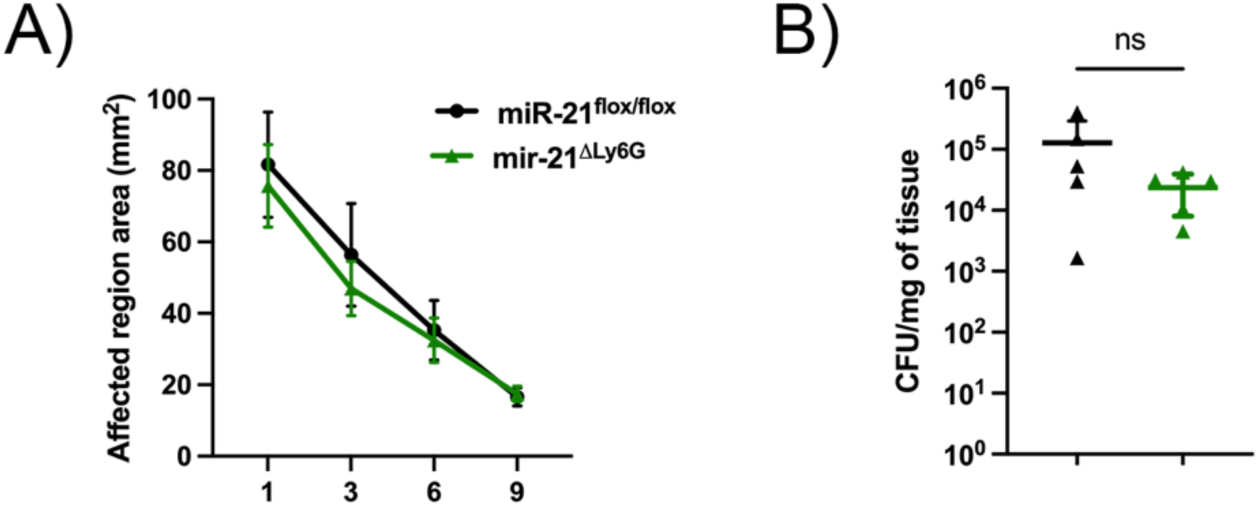
Neutrophil miR-21 deficiency does not alter host defense during *S. aureus* skin infection. miR21^ΔLy6G^ or miR21^flox/flox^ mice were infected with MRSA and monitored throughout the course of infection. **A)** Skin lesion size measured every other day for 9 days post-infection. **B)** Bacterial burden in infected skin quantified by CFU at 9 days post-infection. Data are presented as mean ± SD for five mice from a representative of two independent experiments. *p<0.05 *vs* miR21^flox/flox^ mice.

**Fig. Supp. 2.**
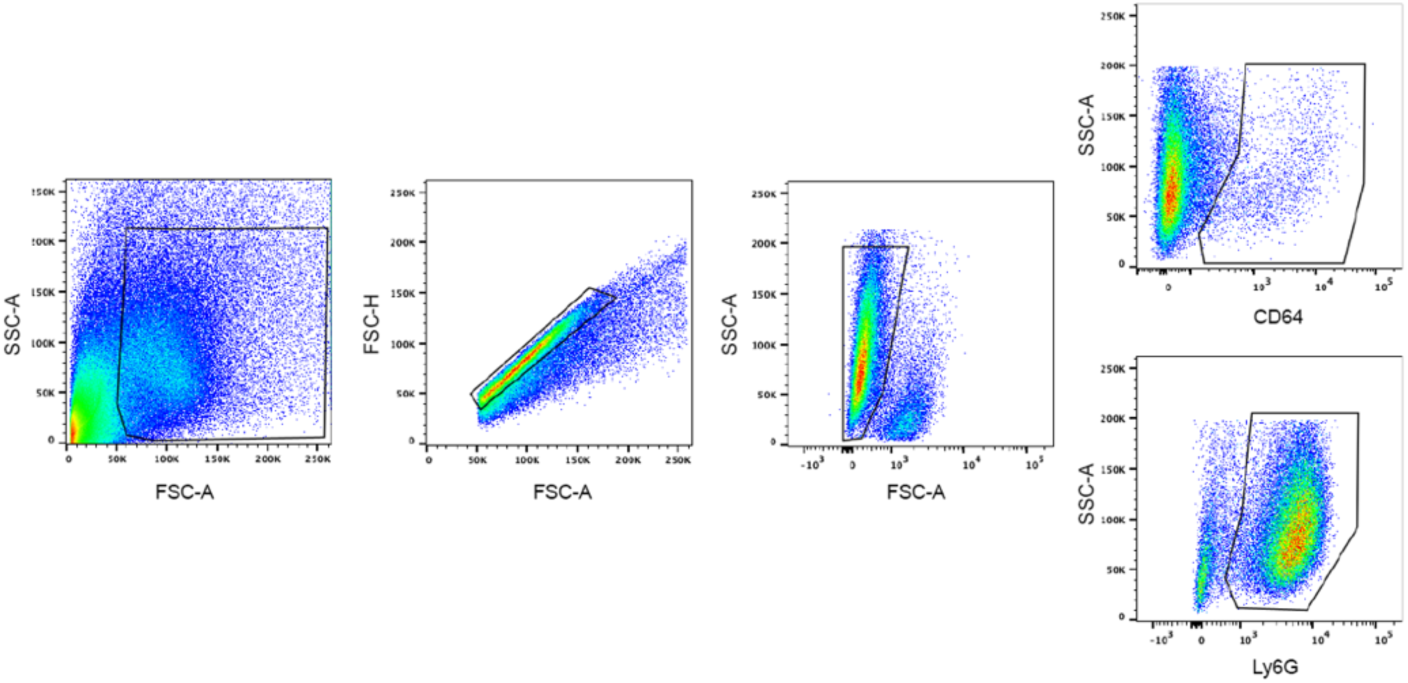
Gate strategy. miR21^Δmyel^ or miR21^flox/flox^ mice were infected with MRSA, and the biopsies were collected at 24 h post-infection. Skin tissue was digested into a single-cell suspension, as described in Materials and Methods, and the total immune cells were characterized by flow cytometry using the following gating strategy: SSC-A x FSC-A/ FSC-H x FSC-A/ SSC-A x Viability/ SSC-A x CD64 or SSC-A x Ly6G. Representative data are shown.

**Fig. Supp. 3.**
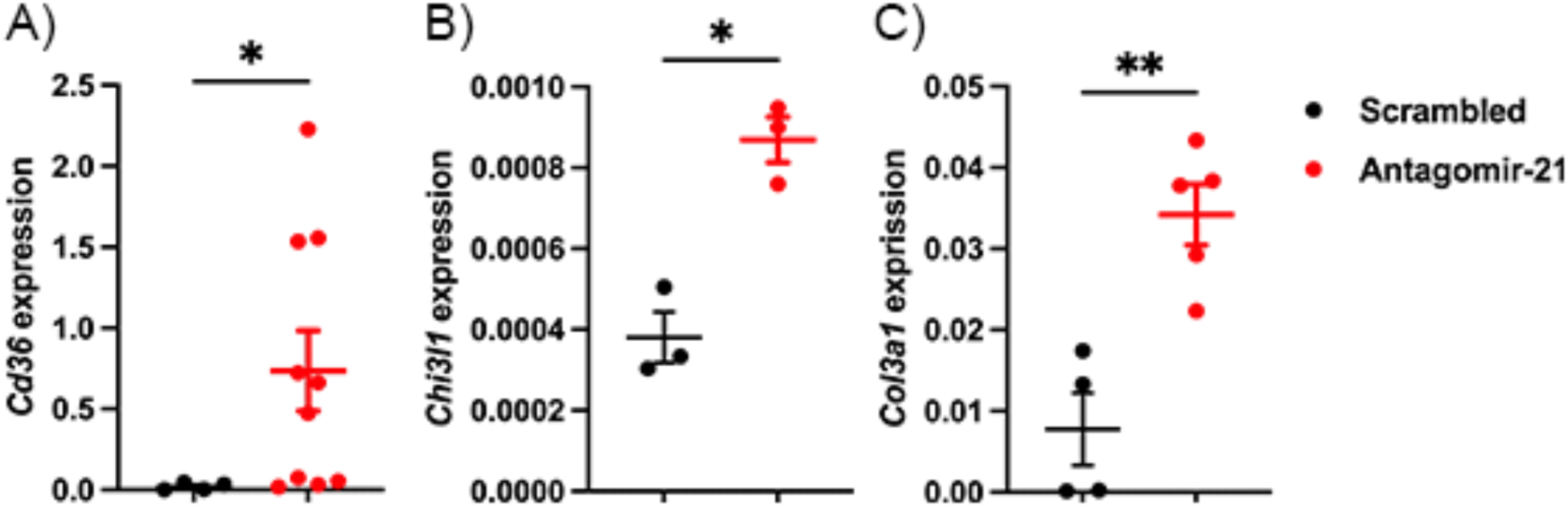
Topical antagomir-21 treatment induces a pro-resolution profile in the skin during *S. aureus* infection. WT mice were infected and treated daily with topical antagomir-21 or scrambled. RNA expression in the skin biopsies was determined by RT-qPCR. **A)** *Col3a1,* **B)** *Chi3l1,* and **C)** *Cd36*. Data are presented as mean ± SD from at least two independent experiments. *p<0.05 and **p<0.005 *vs* scrambled treated mice.

## References

Abtin, Arby, Rohit Jain, Andrew J. Mitchell, et al. 2014. “Perivascular Macrophages Mediate Neutrophil Recruitment during Bacterial Skin Infection.” Nature Immunology 15 (1): 45–53. 10.1038/ni.2769.

Aldrich, Amy, and Tammy Kielian. 2011. “Central Nervous System Fibrosis Is Associated with Fibrocyte-like Infiltrates.” The American Journal of Pathology 179 (6): 2952–62. 10.1016/j.ajpath.2011.08.036.

Ali Alghamdi, Bandar, Intisar Al-Johani, Jawhra M. Al-Shamrani, et al. 2023. “Antimicrobial Resistance in Methicillin-Resistant Staphylococcus Aureus.” Saudi Journal of Biological Sciences 30 (4): 103604. 10.1016/j.sjbs.2023.103604.

Aufiero, Barbara, Meng Guo, Chen Young, et al. 2007. “Staphylococcus Aureus Induces the Expression of Tumor Necrosis Factor-Alpha in Primary Human Keratinocytes.” International Journal of Dermatology 46 (7): 687–94. 10.1111/j.1365-4632.2007.03161.x.

Becker, Russell E. N., Bryan J. Berube, Georgia R. Sampedro, Andrea C. DeDent, and Juliane Bubeck Wardenburg. 2014. “Tissue-Specific Patterning of Host Innate Immune Responses by Staphylococcus Aureus α-Toxin.” Journal of Innate Immunity 6 (5): 619–31. 10.1159/000360006.

Bitschar, Katharina, Christiane Wolz, Bernhard Krismer, Andreas Peschel, and Birgit Schittek. 2017. “Keratinocytes as Sensors and Central Players in the Immune Defense against Staphylococcus Aureus in the Skin.” Journal of Dermatological Science 87 (3): 215–20. 10.1016/j.jdermsci.2017.06.003.

Brandt, Stephanie L., Nathan Klopfenstein, Soujuan Wang, et al. 2018. “Macrophage-Derived LTB4 Promotes Abscess Formation and Clearance of Staphylococcus Aureus Skin Infection in Mice.” PLoS Pathogens 14 (8): e1007244. 10.1371/journal.ppat.1007244.

Brandt, Stephanie L., Nicole E. Putnam, James E. Cassat, and C. Henrique Serezani. 2018a. “Innate Immunity to S. Aureus: Evolving Paradigms in Soft Tissue and Invasive Infections.” Journal of Immunology (Baltimore, Md. : 1950) 200 (12): 3871–80. 10.4049/jimmunol.1701574.

Brandt, Stephanie L., Nicole E. Putnam, James E. Cassat, and C. Henrique Serezani. 2018b. “Innate Immunity to S. Aureus: Evolving Paradigms in Soft Tissue and Invasive Infections.” Journal of Immunology (Baltimore, Md. : 1950) 200 (12): 3871–80. 10.4049/jimmunol.1701574.

Canfrán-Duque, Alberto, Noemi Rotllan, Xinbo Zhang, et al. 2017. “Macrophage Deficiency of miR-21 Promotes Apoptosis, Plaque Necrosis, and Vascular Inflammation during Atherogenesis.” EMBO Molecular Medicine 9 (9): 1244–62. 10.15252/emmm.201607492.

Cheng, Alice G., Hwan Keun Kim, Monica L. Burts, Thomas Krausz, Olaf Schneewind, and Dominique M. Missiakas. 2009. “Genetic Requirements for Staphylococcus Aureus Abscess Formation and Persistence in Host Tissues.” FASEB Journal: Official Publication of the Federation of American Societies for Experimental Biology 23 (10): 3393–404. 10.1096/fj.09-135467.

Cho, John S., Yi Guo, Romela Irene Ramos, et al. 2012. “Neutrophil-Derived IL-1β Is Sufficient for Abscess Formation in Immunity against Staphylococcus Aureus in Mice.” PLoS Pathogens 8 (11): e1003047. 10.1371/journal.ppat.1003047.

Cho, John S., Eric M. Pietras, Nairy C. Garcia, et al. 2010. “IL-17 Is Essential for Host Defense against Cutaneous Staphylococcus Aureus Infection in Mice.” The Journal of Clinical Investigation 120 (5): 1762–73. 10.1172/JCI40891.

Dai, Yuxuan, and Yu Chen. n.d. “Targeting Persistently Activated Inflammatory Microenvironment to Promote Chronic Wound Healing.” Frontiers in Immunology 16: 1708358. 10.3389/fimmu.2025.1708358.

De Melo, Paulo, Annie Rocio Pineros Alvarez, Xiang Ye, et al. 2021. “Macrophage-Derived MicroRNA-21 Drives Overwhelming Glycolytic and Inflammatory Response during Sepsis via Repression of the PGE2/IL-10 Axis.” Journal of Immunology (Baltimore, Md. : 1950) 207 (3): 902–12. 10.4049/jimmunol.2001251.

Deguine, Jacques, and Gregory M. Barton. 2014. “MyD88: A Central Player in Innate Immune Signaling.” F1000Prime Reports 6 (November): 97. 10.12703/P6-97.

Dejani, Naiara N., Stephanie L. Brandt, Annie Piñeros, et al. 2016. “Topical Prostaglandin E Analog Restores Defective Dendritic Cell–Mediated Th17 Host Defense Against Methicillin-Resistant Staphylococcus Aureus in the Skin of Diabetic Mice.” Diabetes 65 (12): 3718–29. 10.2337/db16-0565.

Drury, Ruth E., Daniel O’Connor, and Andrew J. Pollard. 2017. “The Clinical Application of MicroRNAs in Infectious Disease.” Frontiers in Immunology 8: 1182. 10.3389/fimmu.2017.01182.

Feng, Jun, Antai Li, Jingyuan Deng, et al. 2014. “miR-21 Attenuates Lipopolysaccharide-Induced Lipid Accumulation and Inflammatory Response: Potential Role in Cerebrovascular Disease.” Lipids in Health and Disease 13 (February): 27. 10.1186/1476-511X-13-27.

Friedman, Robin C., Kyle Kai-How Farh, Christopher B. Burge, and David P. Bartel. 2009. “Most Mammalian mRNAs Are Conserved Targets of microRNAs.” Genome Research 19 (1): 92–105. 10.1101/gr.082701.108.

Greenlee-Wacker, Mallary C., Kevin M. Rigby, Scott D. Kobayashi, Adeline R. Porter, Frank R. DeLeo, and William M. Nauseef. 2014. “Phagocytosis of Staphylococcus Aureus by Human Neutrophils Prevents Macrophage Efferocytosis and Induces Programmed Necrosis.” The Journal of Immunology 192 (10): 4709–17. 10.4049/jimmunol.1302692.

Hackett, Emer E., Hugo Charles-Messance, Seónadh M. O’Leary, et al. 2020. “Mycobacterium Tuberculosis Limits Host Glycolysis and IL-1β by Restriction of PFK-M via MicroRNA-21.” Cell Reports 30 (1): 124–136.e4. 10.1016/j.celrep.2019.12.015.

Jingjing, Zhang, Zhang Nan, Wu Wei, et al. 2017. “MicroRNA-24 Modulates Staphylococcus Aureus-Induced Macrophage Polarization by Suppressing CHI3L1.” Inflammation 40 (3): 995–1005. 10.1007/s10753-017-0543-3.

Johnston, Daniel G. W., Jay Kearney, Zbigniew Zasłona, Michelle A. Williams, Luke A. J. O’Neill, and Sinéad C. Corr. 2017. “MicroRNA-21 Limits Uptake of Listeria Monocytogenes by Macrophages to Reduce the Intracellular Niche and Control Infection.” Frontiers in Cellular and Infection Microbiology 7 (May): 201. 10.3389/fcimb.2017.00201.

Kawai, Taro, and Shizuo Akira. 2010. “The Role of Pattern-Recognition Receptors in Innate Immunity: Update on Toll-like Receptors.” Nature Immunology 11 (5): 373–84. 10.1038/ni.1863.

Kelly, Alanna M., Emilio G. Vozza, Brenda Morris, et al. 2025. “Staphylococcus Aureus Induces miR-21 Expression to Promote Bacterial Persistence during Nasal Colonisation.” Clinical Immunology (Orlando, Fla.) 281 (August): 110593. 10.1016/j.clim.2025.110593.

Kerckhove, Maiko de, Katsuya Tanaka, Takahiro Umehara, et al. 2018. “Targeting miR-223 in Neutrophils Enhances the Clearance of Staphylococcus Aureus in Infected Wounds.” EMBO Molecular Medicine 10 (10): e9024. 10.15252/emmm.201809024.

Kern, Winfried V. 2010. “Management of Staphylococcus Aureus Bacteremia and Endocarditis: Progresses and Challenges.” Current Opinion in Infectious Diseases 23 (4): 346–58. 10.1097/QCO.0b013e32833bcc8a.

Kertesz, Michael, Nicola Iovino, Ulrich Unnerstall, Ulrike Gaul, and Eran Segal. 2007. “The Role of Site Accessibility in microRNA Target Recognition.” Nature Genetics 39 (10): 1278–84. 10.1038/ng2135.

Kielian, Tammy, Edward D. Bearden, Aaron C. Baldwin, and Nilufer Esen. 2004. “IL-1 and TNF-Alpha Play a Pivotal Role in the Host Immune Response in a Mouse Model of Staphylococcus Aureus-Induced Experimental Brain Abscess.” Journal of Neuropathology and Experimental Neurology 63 (4): 381–96. 10.1093/jnen/63.4.381.

Kluytmans, J., A. van Belkum, and H. Verbrugh. 1997. “Nasal Carriage of Staphylococcus Aureus: Epidemiology, Underlying Mechanisms, and Associated Risks.” Clinical Microbiology Reviews 10 (3): 505–20. 10.1128/CMR.10.3.505.

Kobayashi, Scott D., Natalia Malachowa, and Frank R. DeLeo. 2015a. “Pathogenesis of Staphylococcus Aureus Abscesses.” The American Journal of Pathology 185 (6): 1518–27. 10.1016/j.ajpath.2014.11.030.

Kobayashi, Scott D., Natalia Malachowa, and Frank R. DeLeo. 2015b. “Pathogenesis of Staphylococcus Aureus Abscesses.” The American Journal of Pathology 185 (6): 1518–27. 10.1016/j.ajpath.2014.11.030.

Kong, Eric F., Jennifer K. Johnson, and Mary Ann Jabra-Rizk. 2016. “Community-Associated Methicillin-Resistant Staphylococcus Aureus: An Enemy amidst Us.” PLoS Pathogens 12 (10): e1005837. 10.1371/journal.ppat.1005837.

König, B., G. Prévost, Y. Piémont, and W. König. 1995. “Effects of Staphylococcus Aureus Leukocidins on Inflammatory Mediator Release from Human Granulocytes.” The Journal of Infectious Diseases 171 (3): 607–13. 10.1093/infdis/171.3.607.

Li, Hui-Chun, Chee-Hing Yang, and Shih-Yen Lo. 2022. “Roles of microRNAs in Hepatitis C Virus Replication and Pathogenesis.” Viruses 14 (8): 1776. 10.3390/v14081776.

Li, Yuan, Silei Sui, and Ajay Goel. 2024. “Extracellular Vesicles Associated microRNAs: Their Biology and Clinical Significance as Biomarkers in Gastrointestinal Cancers.” Seminars in Cancer Biology 99 (February): 5–23. 10.1016/j.semcancer.2024.02.001.

Li, Zhouqiang, Hualing Zhang, Zeshan Chen, Guanzhu Wu, Weixing Guo, and Yun Li. 2025. “MicroRNA-21: A Potential Therapeutic Target in Lung Cancer (Review).” International Journal of Oncology 67 (2): 67. 10.3892/ijo.2025.5773.

Liu, Chao, Wei Ouyang, Jingyan Xia, Xiaoru Sun, Liying Zhao, and Feng Xu. 2018. “Tumor Necrosis Factor-α Is Required for Mast Cell-Mediated Host Immunity Against Cutaneous Staphylococcus Aureus Infection.” The Journal of Infectious Diseases 218 (1): 64–74. 10.1093/infdis/jiy149.

Liu, Philip T., Matthew Wheelwright, Rosane Teles, et al. 2012. “MicroRNA-21 Targets the Vitamin D-Dependent Antimicrobial Pathway in Leprosy.” Nature Medicine 18 (2): 267–73. 10.1038/nm.2584.

Liu, Yan-Cun, Xian-Biao Zou, Yan-Fen Chai, and Yong-Ming Yao. 2014. “Macrophage Polarization in Inflammatory Diseases.” International Journal of Biological Sciences 10 (5): 520–29. 10.7150/ijbs.8879.

Luoreng, Zhuo-Ma, Xing-Ping Wang, Chu-Gang Mei, and Lin-Sen Zan. 2018. “Comparison of microRNA Profiles between Bovine Mammary Glands Infected with Staphylococcus Aureus and Escherichia Coli.” International Journal of Biological Sciences 14 (1): 87–99. 10.7150/ijbs.22498.

Ma, Xiaodong, Lindsey E. Becker Buscaglia, Juanita R. Barker, and Yong Li. 2011. “MicroRNAs in NF-kappaB Signaling.” Journal of Molecular Cell Biology 3 (3): 159–66. 10.1093/jmcb/mjr007.

Mehta, Arnav, and David Baltimore. 2016. “MicroRNAs as Regulatory Elements in Immune System Logic.” Nature Reviews Immunology 16 (5): 279–94. 10.1038/nri.2016.40.

Miller, Lloyd S. 2008. “Toll-like Receptors in Skin.” Advances in Dermatology 24: 71–87. 10.1016/j.yadr.2008.09.004.

Miller, Lloyd S., Ryan M. O’Connell, Miguel A. Gutierrez, et al. 2006. “MyD88 Mediates Neutrophil Recruitment Initiated by IL-1R but Not TLR2 Activation in Immunity against Staphylococcus Aureus.” Immunity 24 (1): 79–91. 10.1016/j.immuni.2005.11.011.

Miller, Lloyd S., Eric M. Pietras, Lawrence H. Uricchio, et al. 2007. “Inflammasome-Mediated Production of IL-1β Is Required for Neutrophil Recruitment against Staphylococcus Aureus In Vivo1.” The Journal of Immunology 179 (10): 6933–42. 10.4049/jimmunol.179.10.6933.

Minegishi, Yoshiyuki, Masako Saito, Masayuki Nagasawa, et al. 2009. “Molecular Explanation for the Contradiction between Systemic Th17 Defect and Localized Bacterial Infection in Hyper-IgE Syndrome.” The Journal of Experimental Medicine 206 (6): 1291–301. 10.1084/jem.20082767.

Novelli, M., P. Savoia, I. Cambieri, et al. 2000. “Collagenase Digestion and Mechanical Disaggregation as a Method to Extract and Immunophenotype Tumour Lymphocytes in Cutaneous T-Cell Lymphomas.” Clinical and Experimental Dermatology 25 (5): 423–31. 10.1046/j.1365-2230.2000.00680.x.

Olaru, Florina, and Liselotte E. Jensen. 2010. “Staphylococcus Aureus Stimulates Neutrophil Targeting Chemokine Expression in Keratinocytes through an Autocrine IL-1α Signaling Loop.” Journal of Investigative Dermatology 130 (7): 1866–76. 10.1038/jid.2010.37.

Pasparakis, Manolis, Ingo Haase, and Frank O. Nestle. 2014. “Mechanisms Regulating Skin Immunity and Inflammation.” Nature Reviews Immunology 14 (5): 289–301. 10.1038/nri3646.

Plaut, Roger D., Christopher P. Mocca, Ranjani Prabhakara, Tod J. Merkel, and Scott Stibitz. 2013. “Stably Luminescent Staphylococcus Aureus Clinical Strains for Use in Bioluminescent Imaging.” PLOS ONE 8 (3): e59232. 10.1371/journal.pone.0059232.

Puel, Anne, Capucine Picard, Mathie Lorrot, et al. 2008. “Recurrent Staphylococcal Cellulitis and Subcutaneous Abscesses in a Child with Autoantibodies against IL-6.” Journal of Immunology (Baltimore, Md. : 1950) 180 (1): 647–54. 10.4049/jimmunol.180.1.647.

Putnam, Nicole E., Laura E. Fulbright, Jacob M. Curry, et al. 2019. “MyD88 and IL-1R Signaling Drive Antibacterial Immunity and Osteoclast-Driven Bone Loss during Staphylococcus Aureus Osteomyelitis.” PLoS Pathogens 15 (4): e1007744. 10.1371/journal.ppat.1007744.

Ramirez, Horacio A., Irena Pastar, Ivan Jozic, et al. 2018. “Staphylococcus Aureus Triggers Induction of miR-15B-5P to Diminish DNA Repair and Deregulate Inflammatory Response in Diabetic Foot Ulcers.” The Journal of Investigative Dermatology 138 (5): 1187–96. 10.1016/j.jid.2017.11.038.

Rezaeepoor, Mahsa, Fariba Keramat, Sanaz Jourghasemi, et al. 2024. “MicroRNA -21 Expression as an Auxiliary Diagnostic Biomarker of Acute Brucellosis.” Molecular Biology Reports 51 (1): 264. 10.1007/s11033-023-09193-8.

Riaz, Mahrukh, Muhammad Zohaib Iqbal, Agnes S. Klar, and Thomas Biedermann. 2025. “Immunomodulatory Mechanisms of Chronic Wound Healing: Translational and Clinical Relevance.” MedComm 6 (11): e70378. 10.1002/mco2.70378.

Rodríguez-Morales, Patricia, and Ruth A. Franklin. 2023. “Macrophage Phenotypes and Functions: Resolving Inflammation and Restoring Homeostasis.” Trends in Immunology 44 (12): 986–98. 10.1016/j.it.2023.10.004.

Roy, Sashwati, Suman Santra, Amitava Das, et al. 2020. “Staphylococcus Aureus Biofilm Infection Compromises Wound Healing by Causing Deficiencies in Granulation Tissue Collagen.” Annals of Surgery 271 (6): 1174–85. 10.1097/SLA.0000000000003053.

Schreiber, Heidi A., Jakob Loschko, Roos A. Karssemeijer, et al. 2013. “Intestinal Monocytes and Macrophages Are Required for T Cell Polarization in Response to Citrobacter Rodentium.” The Journal of Experimental Medicine 210 (10): 2025–39. 10.1084/jem.20130903.

Scuruchi, Michele, Angela Avenoso, Federica Aliquò, et al. 2024. “miR-21 Attenuated Inflammation Targeting MyD88 in Human Chondrocytes Stimulated with Hyaluronan Oligosaccharides.” Archives of Biochemistry and Biophysics 759 (September): 110112. 10.1016/j.abb.2024.110112.

Sheedy, Frederick J., Eva Palsson-McDermott, Elizabeth J. Hennessy, et al. 2010. “Negative Regulation of TLR4 via Targeting of the Proinflammatory Tumor Suppressor PDCD4 by the microRNA miR-21.” Nature Immunology 11 (2): 141–47. 10.1038/ni.1828.

Soares, Caroline Leal Rodrigues, Polrat Wilairatana, Larissa Rodrigues Silva, et al. 2023. “Biochemical Aspects of the Inflammatory Process: A Narrative Review.” Biomedicine & Pharmacotherapy 168 (December): 115764. 10.1016/j.biopha.2023.115764.

Stenzel, Werner, Julia Dahm, Monica Sanchez-Ruiz, et al. 2005. “Regulation of the Inflammatory Response to Staphylococcus Aureus-Induced Brain Abscess by Interleukin-10.” Journal of Neuropathology and Experimental Neurology 64 (12): 1046–57. 10.1097/01.jnen.0000189836.48704.ca.

Takeuchi, Osamu, Kiyoshi Takeda, Katsuaki Hoshino, Osamu Adachi, Tomohiko Ogawa, and Shizuo Akira. 2000. “Cellular Responses to Bacterial Cell Wall Components Are Mediated through MyD88-Dependent Signaling Cascades.” International Immunology 12 (1): 113–17. 10.1093/intimm/12.1.113.

Tanaka, Katsuya, Sang Eun Kim, Hiroki Yano, et al. 2017. “*MiR-142* Is Required for *Staphylococcus Aureus* Clearance at Skin Wound Sites via Small GTPase-Mediated Regulation of the Neutrophil Actin Cytoskeleton.” Journal of Investigative Dermatology 137 (4): 931–40. 10.1016/j.jid.2016.11.018.

Tong, Steven Y. C., Joshua S. Davis, Emily Eichenberger, Thomas L. Holland, and Vance G. Fowler. 2015. “Staphylococcus Aureus Infections: Epidemiology, Pathophysiology, Clinical Manifestations, and Management.” Clinical Microbiology Reviews 28 (3): 603–61. 10.1128/CMR.00134-14.

Varikuti, Sanjay, Chaitenya Verma, Erin Holcomb, et al. 2021. “MicroRNA-21 Deficiency Promotes the Early Th1 Immune Response and Resistance toward Visceral Leishmaniasis.” Journal of Immunology (Baltimore, Md. : 1950) 207 (5): 1322–32. 10.4049/jimmunol.2001099.

Wang, Zhuo, Stephanie Brandt, Alexandra Medeiros, et al. 2015. “MicroRNA 21 Is a Homeostatic Regulator of Macrophage Polarization and Prevents Prostaglandin E2-Mediated M2 Generation.” PloS One 10 (2): e0115855. 10.1371/journal.pone.0115855.

Zhang, Fengyuan, Yidan Xia, Jiayang Su, et al. 2024. “Neutrophil Diversity and Function in Health and Disease.” Signal Transduction and Targeted Therapy 9 (1): 343. 10.1038/s41392-024-02049-y.

